# Live imaging human embryos reveals mitotic errors and lineage specification prior to implantation

**DOI:** 10.1101/2024.09.26.614906

**Authors:** Ahmed Abdelbaki, Afshan McCarthy, Anita Karsa, Leila Muresan, Kay Elder, Athanasios Papathanasiou, Phil Snell, Leila Christie, Martin Wilding, Benjamin J. Steventon, Kathy K. Niakan

## Abstract

Meiotic and mitotic chromosome segregation errors in human development have been studied mostly prior to and at the time of fertilisation. Despite chromosomal errors being a leading cause of miscarriage and infertility, chromosome missegregation has not been extensively studied at later stages of human development. Here we optimised labelling, light-sheet live imaging and semi-quantitative analysis of human embryos and reveal chromosome segregation errors just prior to implantation. We found that human embryos exhibited a number of chromosome missegregation events including multipolar spindle formation, lagging chromosomes, misalignment and chromosome slippage. We found that the majority of lagging chromosomes were passively inherited by one of the daughter cells, instead of reincorporating into the nuclei, suggesting a distinct pattern of micronuclei inheritance. By semi-automated segmentation, we tracked the position of labelled cells in human embryos and observed that while most labelled cells remained segregated to the outside, and therefore restricted to a placental-progenitor fate, there was evidence of a rare cell migration to cells positioned on the inside, which suggests that there may be plasticity. Altogether, we found that mitotic chromosome segregation errors arise just prior to implantation, which has implications for our understanding of biological events that contribute to aneuploidy mosaicism.

## Introduction

Following fertilization, human zygotes undergo a series of cell divisions, forming a blastocyst prior to implantation. The majority of human blastocysts display a mix of both normal euploid and abnormal aneuploid cells^1,2,3,4^. In contrast to aneuploidies of meiotic origin, the occurrence of euploid/aneuploid mosaicism is strongly linked to the mitotic errors after fertilization^3, 4^. Aneuploidy is known to contribute to implantation failure and embryonic arrest^5–7^, although mosaic aneuploidy is thought to be tolerated especially in trophectoderm cells^8, 9,10^.

Numerous studies have indicated that mosaicism frequently arises in 30–70% of cleavage-stage embryos^4, 11^ and approximately 10–30% of blastocysts^12, 13^. Evidence for this comes from observation of spindle and nuclear abnormalities in fixed human embryos at static time points. Although mitotic errors in cleavage-stage embryos have been described, mitotic errors in chromosome segregation arising *de novo* at the blastocyst stage in human preimplantation embryos are less well understood.

Characterisation of these mitotic errors at later stages in human embryos has been limited by challenges in nuclear labelling, live cell imaging and tracking. In recent years, advances in live imaging of mouse embryos have expanded our knowledge of chromosome segregation as well as cell fate specification in early mammalian development^14,15,16,17^. There is comparatively limited understanding of cell divisions and timing of cell fate specification in human blastocysts.

Recent reports have successfully achieved labelling of human embryos through microinjection of mRNA in zygote^15^ or by using live DNA dyes^16^. However, microinjections are not suitable for most human embryos donated for research, as these are often obtained at the blastocyst stage. On the other hand, prolonged incubation with live DNA dyes has been shown to induce DNA damage responses and directly impact mitotic progression^17, 18^. Furthermore, all the previous studies imaged human embryos using scanning confocal microscopy, which is only suitable for short term imaging due to high phototoxicity^19^. In contrast, the light-sheet fluorescence microscopy offers a significant improvement in illumination and detection, which minimizes the extent of light exposure and enables long-term imaging^19, 20^.

Here we systematically tested various nuclear labelling methods and developed an electroporation method to introduce mRNA in human embryos at the blastocyst stage. We used live imaging by light sheet microscopy to demonstrate that chromosome segregation errors arise in human blastocysts including *de novo* multipolar spindle formation, lagging chromosomes, misalignment and chromosome slippage. By tracing the lagging chromosomes that arise following mitosis, we observed that the micronuclei formed were passively inherited by one of the daughter cells and not reincorporated in subsequent mitoses. We developed a semi-automated pipeline to track the position of labelled cells at the blastocyst stage and found that while the majority of cells remain specified to a placental progenitor compartment, there was a rare contribution of an individual cell to the inner cells within the blastocoel cavity. Altogether we optimised a method to characterise in detail a previously challenging to image stage of human development, thereby revealing the origins of mitotic errors and clarifying the timing of lineage specification.

## Results

### Optimised strategies for nuclear DNA labelling in mouse and human embryos

To track and trace nuclei during live imaging of human preimplantation development we initially systematically investigated various labelling methods using mouse embryos. We required a method that allowed us to label embryos with high efficiency and to track cells for 48hrs without negatively influencing cell proliferation or development. We compared fluorescent labelling methods using lentivirus, adeno-associated virus (AAV), baculovirus (BacMam), DNA dyes, and electroporation of mRNAs.

We initially labelled four-cell stage mouse embryos because this allowed us to determine the perdurance of labelling for 48hrs up to the mid-blastocyst stage. Mouse embryos were transduced with either high-titer lentivirus carrying an H2B-GFP reporter, BacMam H2B-GFP, or AAV serotype 6 (AAV6)-GFP. Theses vectors expressed GFP under the control of a constitutive elongation factor 1a (EF1a) promoter. We monitored embryo development and fluorescent expression over time for 48hrs. We showed that while lentiviral transduction in HEK 293T cells is robust, H2B-GFP expression was not detected following lentiviral transduction in any of the embryos analysed (Extended Data Fig. 1a, b, c), indicating silencing. Baculovirus (BacMam) showed faint signals in one cell at morula stage in 1/20 embryos imaged (Extended Data Fig. 1d, e, f). Adeno-associated virus (AAV6) exhibited transient low expression, lasting only 24 hours (Extended Data Fig. 1g, h, i). We therefore excluded these methods from subsequent analysis in this early embryo context.

We next examined different DNA dye labelling methods in live cells including SPY650-DNA, 5-TMR-Hoechst, 4-TMR-Hoechst, 4-580CP-Hoechst, 5-580CP-Hoechst and Nuclight Rapid Red. Mouse embryos were cultured continuously in media containing working concentrations of different DNA dyes from the four-cell stage and imaged after 48hrs. Among all the tested DNA dyes, SPY650-DNA dye stained the majority of cells at the cleavage stage. This initially looked promising however, only nuclei of TE cells at the blastocyst stage were labelled whereas in the ICM there was nonspecific staining of the cytoplasm (Extended Data Fig. 2a, b, c).

To further explore methods for nuclear DNA labelling, we optimised mRNA electroporation of mouse cleavage stage embryos using an *H2B-mCherry* mRNA. We identified electroporation parameters that allowed chromosome labelling following electroporation of mouse embryos at cleavage and blastocyst stages. We observed that electroporation of mRNA from the 4-cell stage at a concentration ranging from 500 to 800 ng/µl had no obvious impact on total cell number, the proportion of trophectoderm or epiblast cells, or progression of development to the blastocyst stage (Extended Data Fig. 3a, b). Following *H2B-mCherry* mRNA electroporation (Extended Data Fig.3c), we quantified the expression of lineage associated molecular markers of the trophectoderm (CDX2) and epiblast (NANOG) (Extended Data Fig. 3d, e). We observed no difference in total cell number between electroporated and control embryos. Additionally, there was no significant difference in the number of CDX2- or NANOG-positive cells (Extended Data Fig. 3f, g, h). We therefore progressed with the use of mRNA electroporation for nuclear DNA labelling of mouse and human embryos prior to live imaging.

### Light sheet live imaging of mitotic timing in mouse and human embryos

We next optimised live imaging of nuclear labelled mouse embryos using light sheet microscopy based on methods published previously^19^, prior to imaging human preimplantation embryos. We selected the LS2 light sheet microscope because it has dual illumination and double detection to capture dual view of samples (Fig. 1a, b, c). Notably, both developmental timing and blastocyst progression were not significantly different between light-sheet imaged and non-imaged control embryos (Fig. 1d, e, f).

**Figure 1.**
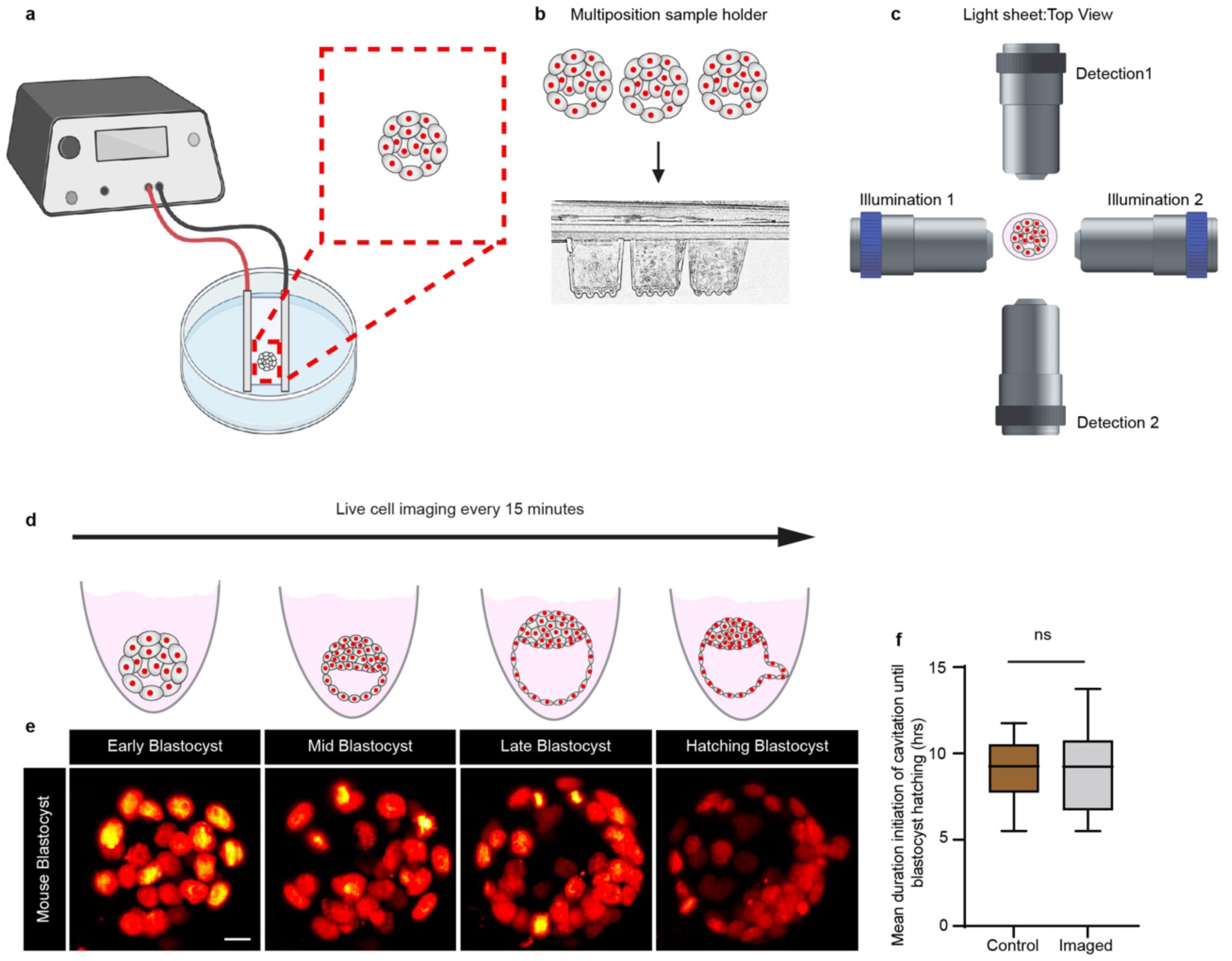
Dual-view light-sheet microscope imaging of preimplantation embryos. **a,** Schematic of the experimental setup: Mouse embryos were electroporated with 500ng/µl H2B-mCherry mRNA and allowed to recover for 2 hours before live-cell light-sheet imaging. **b**, Embryos were transferred to the multi-position sample holder. **c,** Schematic showing the configuration of two detection and two illumination chambers from the microscope’s top view of the sample chamber. **d,** Mouse embryos expressing H2B-mCherry were imaged by light-sheet microscopy from early to late blastocyst stages at the selected time points shown. Scale bar, 20 µm. **f,** The duration of the initiation of cavitation until blastocyst hatching was quantified in mouse embryos labelled with H2B-mCherry and subject to live-cell light-sheet imaging compared to unlabelled control embryos that were imaged using brightfield in light-sheet imaging. Box plot shows median (horizontal black line), mean (small black squares), 25th and 75th percentiles (boxes), 5th and 95th percentiles (whiskers). Both groups exhibit similar developmental timing (n =10 embryos per group: NS, not significant by Student’s t-test).

Light sheet imaging following nuclear DNA labelling with *H2B-mCherry* allowed us to track the phases and duration of mitosis (prophase, metaphase, anaphase and telophase) in mouse embryos (Fig. 2a, b and Supplementary video 1). We subsequently thawed and electroporated early human blastocysts 5 days post fertilisation (dpf) with equivalent concentrations of *H2B-mCherry* mRNA and live imaged them by light sheet microscopy for up to 46hrs (Supplementary video 2). Mitosis was defined as the interval between a prophase and the first signs of telophase, and here we focused on the early to mid-blastocyst stages of mouse and human development (Fig. 2c, d). We determined that there was a similar duration from the start to end of mitosis between the species, which was 49.8 ± 12.7 min in human embryos (n = 73 cells from 7 human embryos) and 47.3 ± 9.6 min in mouse embryos (n = 73 cells from 10 mouse embryos, p = 0.2373) (Fig. 2e). By contrast, we found that interphase was longer in human embryos compared to the mouse (18.73 ± 4.89 versus 11.33 ± 3.49 h, p < 0.0001) (Fig. 2f). This suggests that difference in the timing of interphase is a major contributor for setting the pace of preimplantation development across different species^21^.

**Figure 2.**
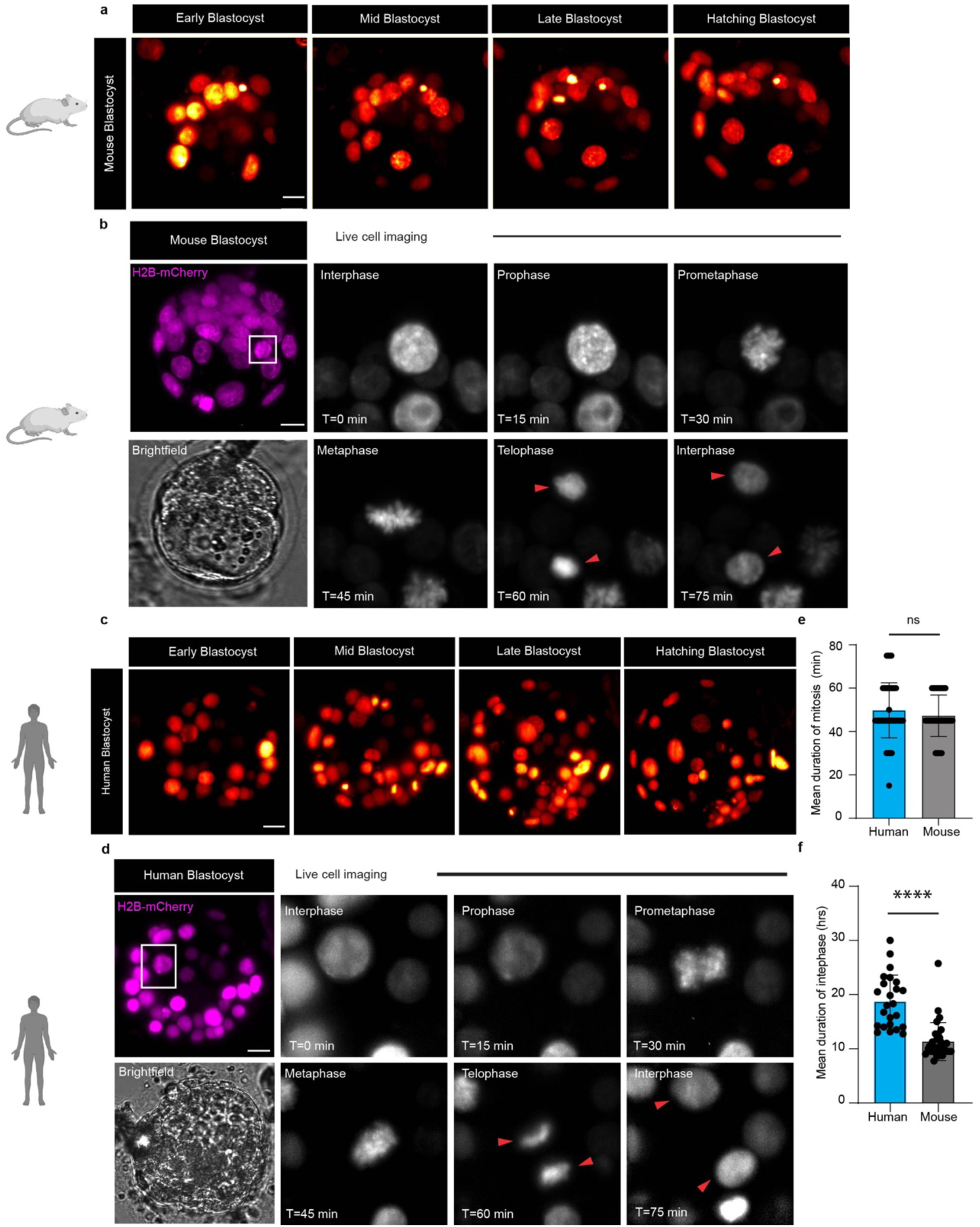
Live-cell time-lapse imaging of mouse and human pre-implantation embryos reveals cell cycle progression and mitosis. **a, c,** Live-cell time-lapse light-sheet imaging of embryos labelled with H2B-mCherry enables the visualization of nuclei as mouse and human embryos progress from early, mid, late, and hatching stages of blastocyst development. **b, d,** Time lapse images of mitosis (prophase, metaphase, anaphase and telophase) in both mouse and human cells at selected time points shown. **e, f,** Quantification of the duration of interphase and mitosis in mouse and human cells at the blastocyst stage (n = 10 mouse and n = 7 human blastocysts; n = 73 mitotic cells and n = 24 interphase cells in mouse blastocysts and n = 73 mitotic cells and n = 30 interphase cells for human blastocysts); ∗∗∗∗p < 0.0001, NS, not significant by Student’s t-test). Error bars represent s.d. Scale bar, 20 µm. See Video S1, S2.

### Characterization of mitotic errors in mouse and human blastocysts

We reasoned that our ability to examine all stages of mitosis would allow us to characterise chromosomal segregation errors in blastocyst stage human embryos. To identify mitotic errors, we analyzed the dynamics of chromosome segregation from 5 to 7 dpf in blastocyst stage human embryos (n = 100 cell divisions across 7 labelled human blastocysts) and compared this to chromosome segregation in mouse embryos at equivalent stages from 3.25 to 4 dpf (n = 285 cell divisions across 15 labelled mouse blastocysts) (Fig. 3a, b). Prior to the onset of anaphase, chromosomes are fully aligned in 95% of mouse blastocyst stage cells, while 89% of chromosomes in human blastocyst stage cells are similarly aligned. Misaligned chromosomes were observed in 9% of human cells compared to 4% in mouse cells (Fig. 3c, d and Extended Data Fig. 4). This included lagging chromosomes and micronuclei formation in one of the daughter cells of either human or mouse blastocyst stage cells (Fig. 3a, b and Supplementary video 3).

**Figure 3.**
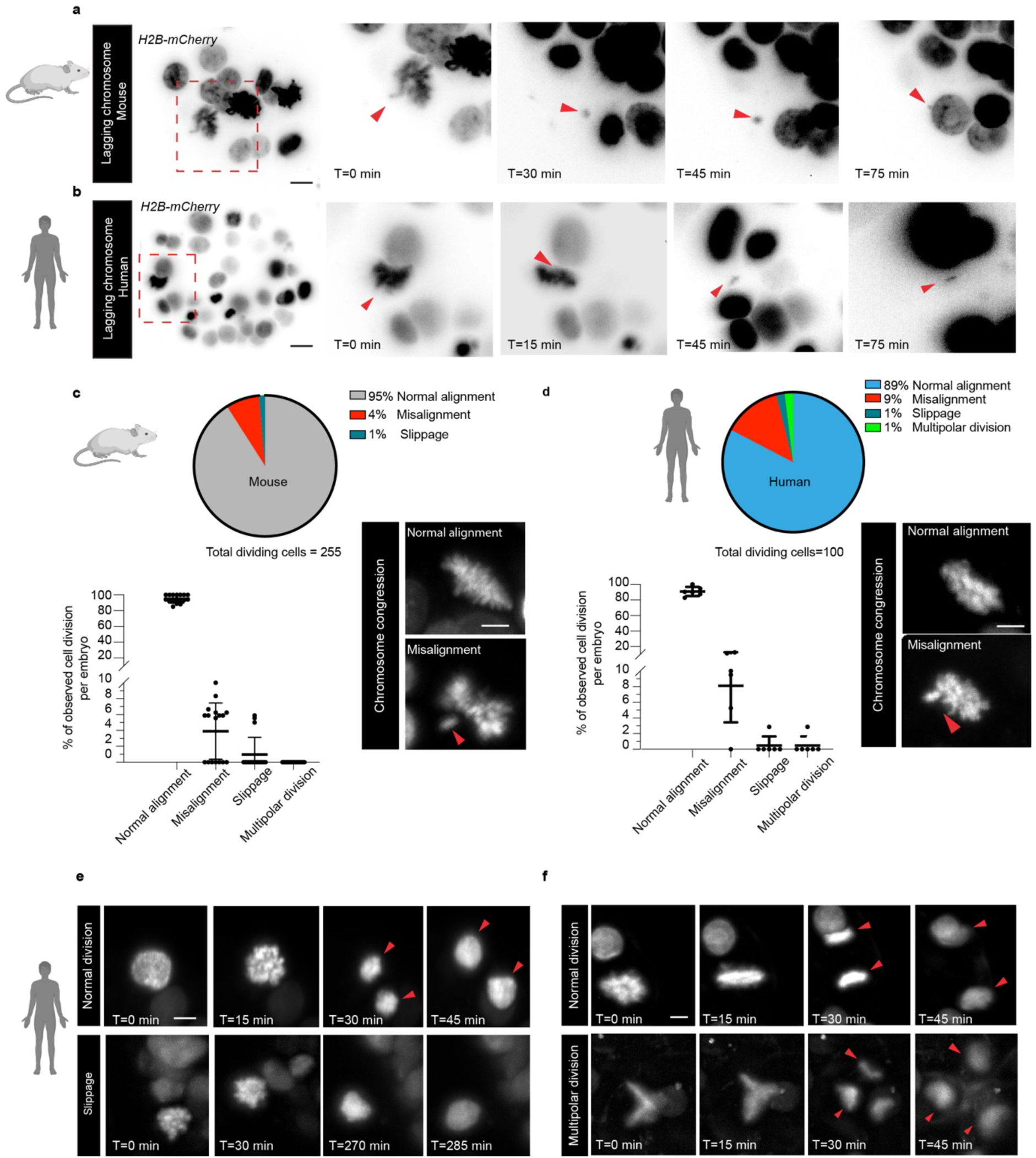
Chromosome segregation errors in mouse and human embryos at the blastocyst stage. **a,b,** Examples of live-cell time-lapse light-sheet imaging of mouse and human embryos expressing H2B-mCherry mRNA enables the identification of misaligned chromosomes, lagging chromosomes and the formation of micronuclei. **c,d,** Analysis of chromosome alignment prior to anaphase and mitotic errors in H2B-mCherry-expressing embryos (n = 17 mouse and n = 6 human embryos, n = 255 and n = 100 mitotic cells for mouse and human, respectively). Error bars represent s.d. Time-lapse imaging illustrating the multipolar division (**e**) and mitotic slippage (**f**) in human embryos at the selected times shown. Daughter cells indicated with arrowheads. Scale bar, 20 µm. See Video S3, S4, S5.

We observed a rare event in 1% of human blastocyst cells, whereby a TE cell prematurely exits from mitosis and, in the absence of chromosome segregation and cell division, enters the next G1 phase of the cell cycle as a tetraploid cell (Fig. 3e and Supplementary video 4). This is a process referred to as mitotic slippage^22, 23^. In mouse embryos slippage was observed in approximately 1% of dividing trophectoderm cells and presumptive tetrapoid cells. We observed in only human embryos some blastocyst stage cells that underwent multipolar cell divisions in 1% of dividing cells. Multipolar spindle formation in anaphase resulted in the production of three daughter cells (Fig. 3f and Supplementary video 5). It is unclear whether the multipolar spindle formation observed in blastocyst stage human cells reflects mechanisms observed in somatic cells that fail to satisfy the spindle assembly checkpoint (SAC) and undergo a mitotic delay known as D-mitosis, whereby delayed mitotic cells either die in mitosis, segregate their chromosomes into two or more aneuploid progeny, or escape mitosis and enter G1^23, 24^. We conclude that misaligned chromosomes, multipolar division, and slippage contributes to mosaic aneuploidy at the blastocyst stage in human embryos.

### The fate of micronuclei in mouse and human embryos

Micronuclei are frequently observed in cancer cells and in mammalian embryos, including humans and mice^25,26,27,28^. While the predominant fate of the micronuclei is persistence in the cytoplasm and passive inheritance during subsequent mitosis in mouse embryos^26^, in cancer cells, chromosomes within micronuclei reintegrate into the nucleus. To understand the fate of micronuclei, we tracked what occurs after their formation in human and mouse blastocysts using light sheet live cell imaging. Surprisingly, the majority of micronuclei (89%) were maintained in the cytoplasm and did not appear to fuse with the nucleus during interphase. Following nuclear envelope breakdown, micronuclei remained separated from the chromosomes of the nucleus throughout the M-phase (Fig. 4a, b). The micronuclei were consistently passively inherited in one of the daughter cells in both mouse and human embryos (Fig. 4c, d). Within the duration of imaging, we also observed that micronuclei reintegrate in 11% of the cases in human blastocysts and 7% of the cases in mouse blastocysts (Extended Data Fig. 5 and Supplementary video 6). Moreover, immunofluorescence analysis of human embryos cultured in conventional incubation conditions shows that micronuclei were also detected in the absence of live-cell light-sheet imaging (Fig. 4e, f). Altogether these finding revealed a unique conserved pattern of micronuclei inheritance in mouse and human embryos.

**Figure 4.**
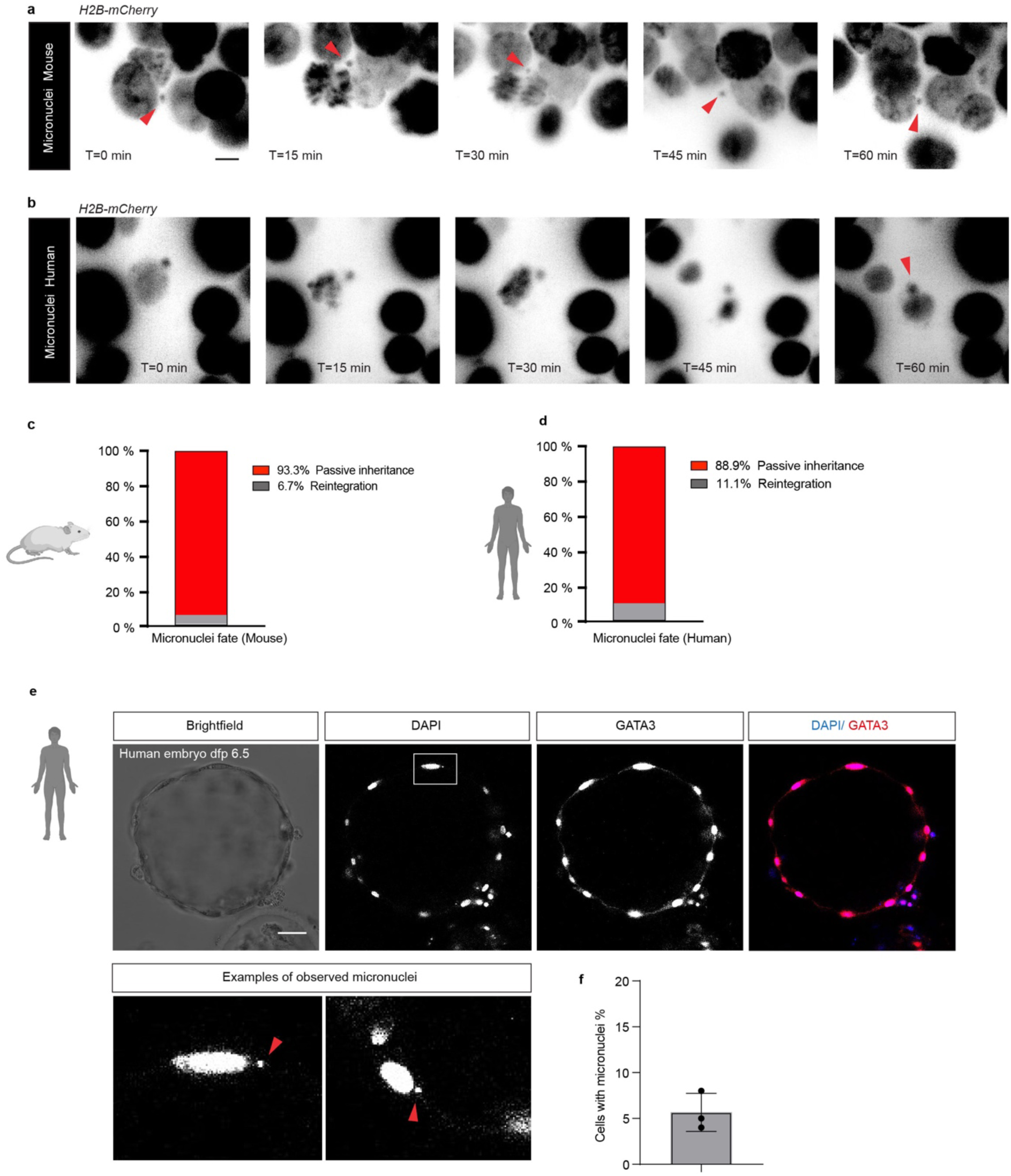
Micronuclei inheritance in mouse and human blastocysts. **a,b,** Example of micronuclei inheritance observed in H2B-mCherry-expressing mouse and human embryos during live-cell time-lapse light-sheet imaging. Micronuclei (red arrow) stays distinct from the rest of the chromosomes throughout the mitosis and was inherited by one of the daughter cells in mouse embryos and human embryos. Inverted images black and white are used for better visualization. **c,d,** Percentage of micronuclei reintegration and passive inheritance in mouse and human embryos. **e**, Human embryo cultured in a conventional incubator and not subject to light-sheet imaging indicating presence of micronuclei. Immunofluorescence analysis of a fixed human blastocyst at 6.5 dpf stained for GATA3 (trophectoderm molecular marker, magenta) and DAPI (blue) nuclear labelling. illustrating the appearance of micronuclei. **f,** Percentage of cells with micronuclei in human blastocysts cultured in a conventional incubator in the absence of light-sheet imaging, Mean ± S.D. of n = 3). Micronuclei indicated with an arrowhead. Error bars represent s.d. Scale bar, 30 µm. See Video S6.

### Trophectoderm cells are largely specified in human blastocysts with a rare contribution to inner cells

The timing of extraembryonic trophectoderm, placental progenitor, specification occurs at the morula stage in 16-cell stage mouse embryos using reporter labelling as well as live-embryo light sheet microscopy, but equivalent studies in human development have not been performed^29,19^. Previous studies labelling human trophectoderm and either reaggregating only labelled cells, or aggregating labelled cells with unlabelled human embryos, suggested that human trophectoderm cells at 5 dpf are not yet irreversibly committed and can contribute to both trophectoderm and to uncommitted inner cell mass cells that give rise to the embryonic epiblast and yolk sac progenitor cells^27^. However, disaggregation/reaggregation studies are not equivalent to lineage tracing studies in unperturbed embryos because the disaggregation technique is known to impact on cell fate.

To track and trace the eventual position of individual labelled nuclei in embryos, we initially developed a semi-automated nuclei segmentation method using a customized version of the StarDist-3D network architecture (Extended Data Fig. 6a). To optimise input image quality, the dual-view light sheet images were averaged for contrast enhancement, data resampled to match lateral and axial resolutions, intensity normalized and finally, a gamma correction was applied. Given the variability in the size and shape of nuclei, especially in human blastocysts, we altered the network architecture to increase the receptive field for segmentation. Training was performed on annotated mouse embryo images. (Extended Data Fig. 6b, c). Nuclei in 3D timelapse images were then automatically segmented, and tracking was performed using regularized Gaussian mixture optimal transport with added regularization to enhance the quality of automated tracking. The 3D segmentation and tracking were validated using Fiji’s TrackMate tool for manual correction. We initially trained the pipeline on annotated mouse embryo images (Extended Data Fig. 6b, c).

To determine whether outer trophectoderm cells are specified, we labelled mouse embryos at the cleavage stage (8-cell stage at 2.5 dpf), followed DNA labelled nuclei in mouse embryos in early (3 dpf) and mid-blastocysts (3.5 dpf) and determined their final position within the blastocysts. Tracking mouse embryo cells showed that cleave stage blastomeres (2.5 dpf) gave rise to daughter cells that contribute to both the trophectoderm and the cells of the inner cell mass. By the early blastocyst stage (3 dpf) cells are specified to either trophectoderm or inner cell mass daughter cells (Extended Data Fig. 6d), consistent with previous studies^28,29,30,31,32^.

We next tracked labelled cells in human embryos (Fig. 5a, b), which showed that the majority of outer trophectoderm cells remained on the outside and gave rise to more trophectoderm cells in intact embryos (Fig.5 c-h, Extended Data Fig. 7 a, b and Supplementary video 7, 8, 9). Surprisingly, in one embryo we observed a single cell at 6.25 dpf which was transiently positioned on the outside of the embryo, overlaying the inner cell mass, and once the cell divided, one daughter cell remained on the outside, while the other cell migrated to the inside position following cell division (Extended Data Fig. 7 a, b and supplementary video 7, 10, 11). No further change in position was observed during the imaging and it was assumed that the cell remained in the inside. After imaging, human embryos were fixed and stained for lineage-associated markers (Extended Data Fig. 7c, d). We detected the expression of molecular markers of the trophectoderm (GATA3) and epiblast (NANOG), indicating that light sheet imaging did not obviously perturb human embryo development (Extended Data Fig. 7d). Moreover, we detected GATA3 expression in cells positioned inside the blastocoel cavity adjacent to the polar trophectoderm. Thus, we observed that it was rare for trophectoderm cells to change their position from the early to mid-blastocyst stage of human development.

**Figure 5.**
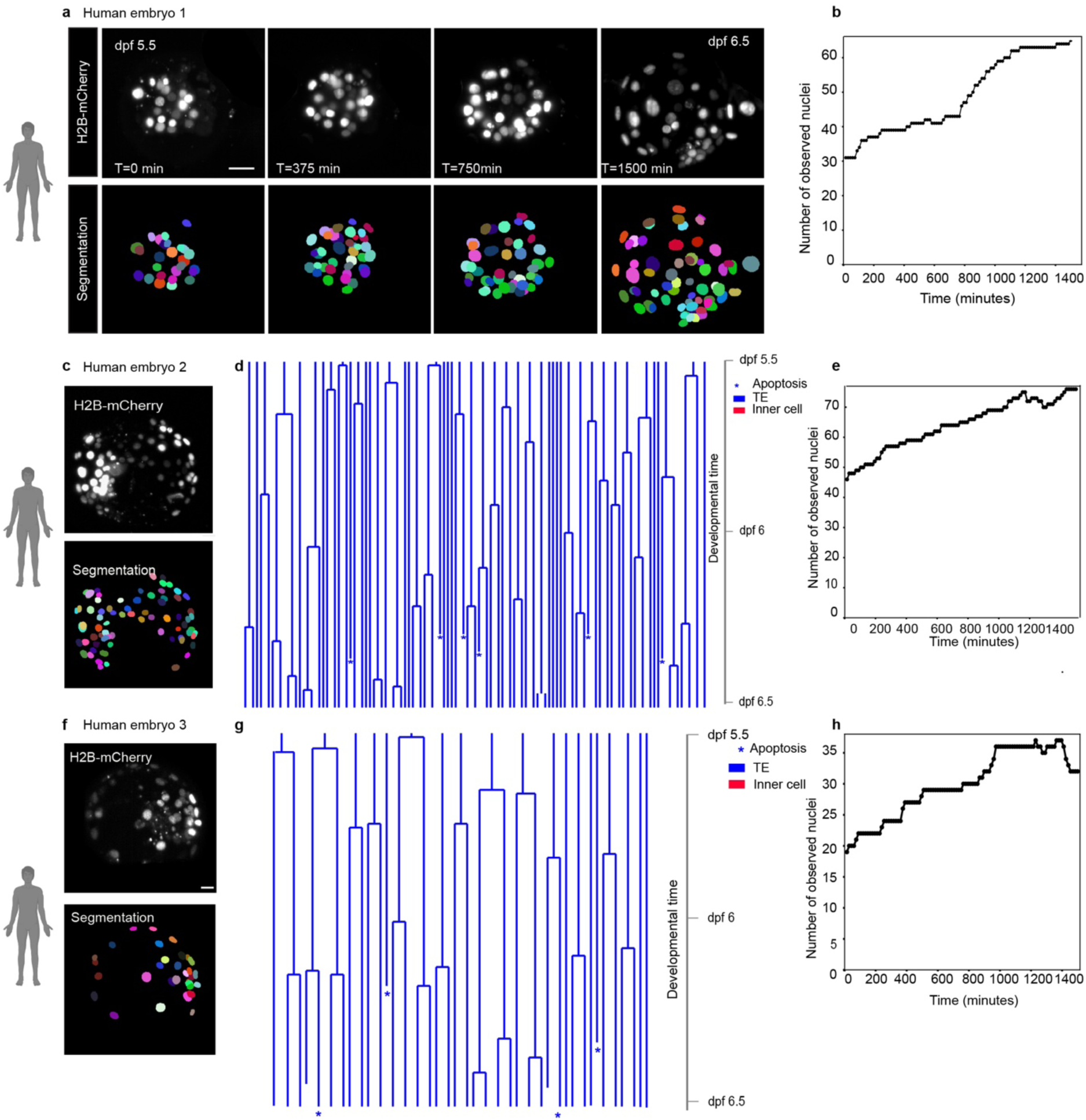
Cell fate specification in human blastocysts **a,** Live-cell time-lapse imaging and 3D segmentation of human embryo at different time points from 5.5 to 6.5 dpf at the selected time points indicated. The fluorescent signals of H2B-mCherry were used to track cells. **c, f,** Selected frames from time-lapse imaging and 3D segmentation of an H2B-mCherry expressing human embryos. **d, g,** Lineage tree of human pre-implantation development from the early to late blastocyst stage is shown. Each H2B-mCherry labelled cell is represented on the tree. Each cell represented on the tree is color coded according to position. **b, e, h,** Number of observed nuclei across time is shown.

We next examined how changes in the size and shape of the human embryo impact on nuclear orientation and division (Fig. 6a). We observed oscillations in the average volume and anisotropy of some of the human embryos over time as they underwent shape changes from a spherical to an elongated oblong blastocyst that hatched from a glycomembrane shell called the zona pellucida (Fig. 6b, c and Extended Data Fig. 8a). Notably, the average nuclear volume and anisotropy exhibited similar dynamic changes over time in the same embryos (Fig. 6d, e and Extended Data Fig. 8b). We quantified nuclear orientation and division with respect to cell position within the embryo and found that the nuclei were oriented tangentially rather than radially (K-S test p<10^−300^) (Fig. 6f, h) in human embryos. Furthermore, an analysis of cell division angle also demonstrated a bias toward tangential division over radial division (K-S test p<10^−9^) (Fig. 6g, i) in human embryos. These findings suggest that nuclear shape and division angle can be linked to tissue geometry.

**Figure 6.**
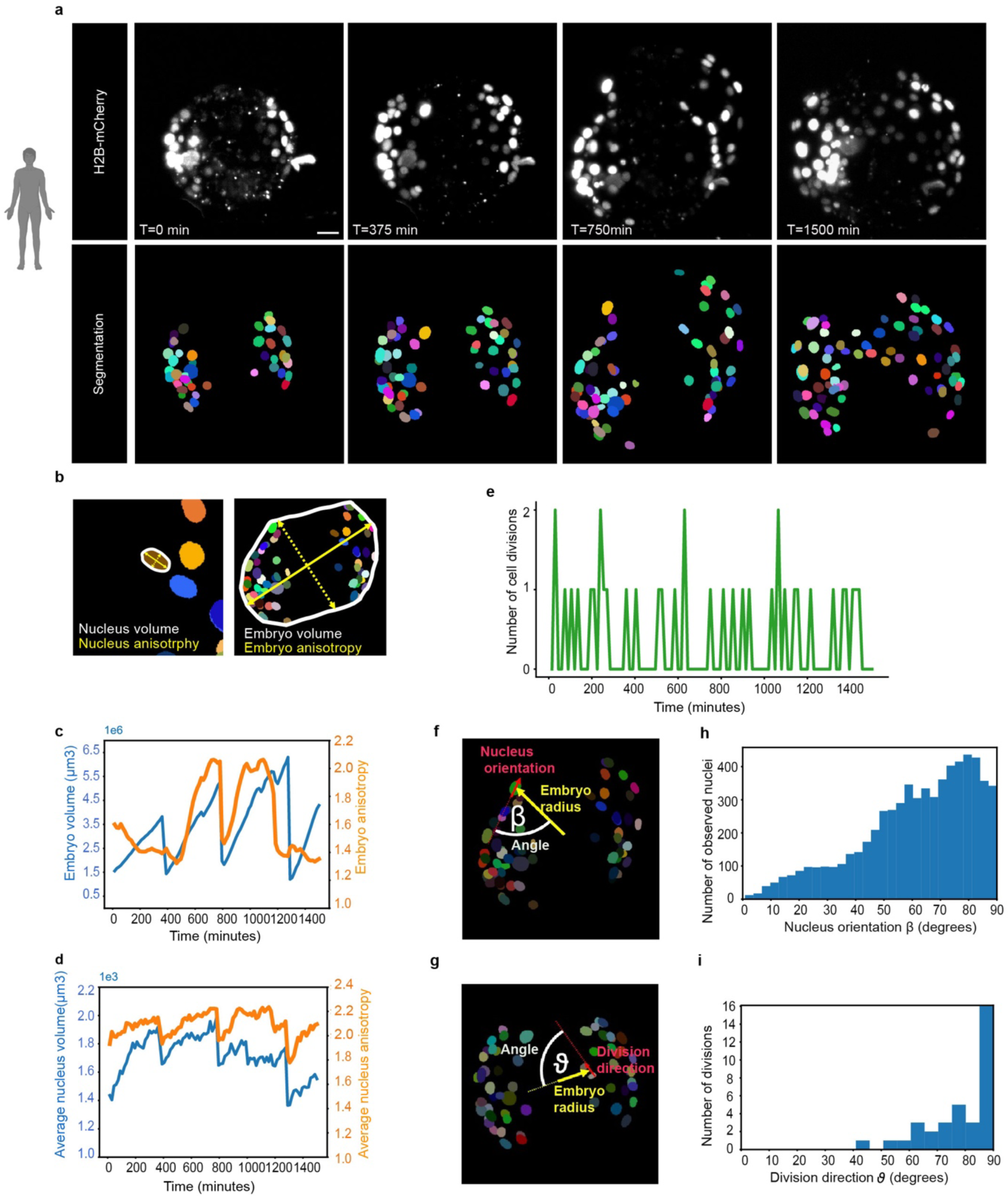
Nuclei shape, size, and division axis orientation in human blastocysts **a,** 3D segmentation of human embryo nuclei at different time points following imaging. **b**, Schematic illustrating how embryo and nuclei size and shape were measured. **c**, Embryo volume (blue) and anisotropy (orange) across time. **d,** Average nucleus volume (blue) and anisotropy (orange) across time. **e,** Number of cell divisions across time. **f,** Schematic illustrating how the angle (β) of the nucleus orientation (red arrow) with respect to the embryo center (yellow arrow) was measured. **g,** Schematics illustrating how the angle (α) between the embryo (yellow arrow) and nucleus (red arrow) orientation was measured. **h,** Histogram of β across all time points showing that nuclei prefer to be oriented tangentially rather than radially to the centre of the embryo (K-S test p<10^−300^). **i,** histogram of ϑ across all time points showing that cells strongly prefer to divide tangentially rather than radially to the centre of the embryo (K-S test p∼10^−12^).

## Discussion

Here we optimised a combination of methods for nuclear labelling and light sheet microscopy to image human embryos up to 46 hrs of development from early to mid-blastocyst stages. This is considerably longer than the 12h reported in previous publications using spinning disk microscopy^15, 16^. Our live imaging of human and mouse embryos provides important insights into mitotic errors, micronuclei formation and inheritance as well as trophectoderm cell fate specification at the blastocyst stage.

We identified species-specific differences in cell cycle length between humans and mice. While the duration of mitosis remains relatively similar, interphase is longer in human embryo cells compared to mice. These findings are consistent with a previous study suggesting that the longer duration of human preimplantation development, compared to the mouse, is attributed to differences in cell cycle length^16^. In the future it would be interesting to further investigate cell cycle progression using reporters of different stages of the cell cycle to have a more accurate timing of mitotic entry and exit^33^.

In addition, our live imaging of human blastocysts revealed mitotic errors, including multipolar chromosome segregation and mitotic slippage. Multipolar chromosome segregation may arise from supernumerary centrosomes, chromosomal instability, or the loss of spindle pole integrity in cells with a normal number of centrosomes^30, 31^. Occasionally, embryonic cells may acquire extra diploid complement of chromosomes (leading to a tetraploid state) in human preimplantation embryos ^32^. The precise origin of tetraploidy remains unclear, but potential explanations include cytokinesis failure and cell fusion^34^. Our data suggest that mitotic slippage also plays a role in driving tetraploidy in human embryos where cells experiencing a delay in mitosis exit without proper chromosome segregation to avoid cell death. We speculate that mitotic slippage may also provide an additional mechanism, besides endoreduplication, for the generation of tetraploid cells in mice^35, 36,37^. Detailed analysis of centrosomes, spindle formation, and the spindle checkpoint throughout human and mouse preimplantation development would inform the underlying causes of these mitotic errors.

Sever misalignment can result in lagging chromosomes which frequently end in micronuclei^25, 26, 38, 39^. Using our less invasive live-cell imaging system, tailored for prolonged imaging of mammalian pre-implantation embryos, we traced the fate of micronuclei in both mouse and human embryos. Unlike cancer cells, our findings indicate that the majority of micronuclei are passively inherited by one of the embryonic daughter cells. This is consistent with previous work from mouse embryos^26^. The fate of cells with micronuclei and whether these cells are eventually eliminated at later stages of development remains an open question. In somatic human cell lines and cancer cells, such chromosomes in micronuclei are subject to defective DNA replication and activate the interferon inflammatory signalling response through recognition by the viral receptor cyclic GMP-AMP synthase (cGAS)^40, 41^. It remains unclear whether these events seen in somatic cells are similar in early mammalian development.

Initiation of the trophectoderm transcriptional program, marked by GATA3 expression in outer cells, occurs at the morula stage and is conserved across species, including the mouse, cow and human embryos^42, 43^. By tracking cells expressing H2B-mCherry in human blastocysts, we observed the rare inward migration / internalization of an outer trophectoderm cell into the inner position, largely consistent with mouse mid-blastocyst trophectoderm cells that remain committed to outside trophectoderm cells^28,29,44,45,46^. Unlike the trophectoderm of mouse blastocysts, trophectoderm cells of human, cow, pig and rabbit blastocysts have longer perdurance of molecular markers of the epiblast including OCT4 and SOX2^47,48,49^. Human blastocyst aggregation assays using trophectoderm cells isolated from 5 dpf human embryos shows that the trophectoderm cells are able to contribute to NANOG-expressing inner cell mass and can also give rise to blastocysts comprised of trophectoderm and inner cell mass^27^, thereby suggesting plasticity at this stage. Additionally, in bovine embryos, morula aggregation assays suggest that trophectoderm cells retain the ability to give rise to the inner cell mass until at least the expanding blastocyst stage^47^. Altogether, this suggests that trophectoderm cells may not yet irreversibly committed up to the mid-blastocyst stage in human embryos, but further labelling and lineage tracing studies are needed to determine the frequency of this, so far, rare observation.

Our findings from live imaging analysis are consistent with a recent study suggesting that trophectoderm cells in the late human blastocyst stage undergo multilayering and contribute to inner cells within the blastocoel cavity^50^. In the future, it will be important to characterise gene and protein expression of internalised cells to determine if they express molecular markers of the trophectoderm, epiblast or yolk sac progenitor cells after live imaging using registration approaches and to track and trace the cells for longer in development. It will be interesting to determine the molecular mechanisms that regulate how human cells undergo cell fate determination and become irreversibly committed in their fate and function. The approaches we developed will be useful to address whether there is indeed bias of cells at the 2-cell stage in human embryos, as has been recently suggested^51^. Altogether we have developed methods for nuclear labelling, tracking and tracing of live human embryos, thereby revealing mechanisms of chromosome mis-segregation and early cell fate decisions. These methods can be used in the future in challenging to study and sensitive developmental contexts and to extend the time of live imaging human embryos to gain further insights into early embryogenesis.

## Materials and Methods

### Ethics statement

This study was approved by the UK Human Fertilisation and Embryology Authority (HFEA): research licence numbers R0162, R0397, R0401 and R0152 and independently reviewed by the Health Research Authority’s Research Ethics Committee IRAS projects 308099, 252286 and 272218.

The process of licence approval entailed independent peer review along with consideration by the HFEA Licence and Executive Committees and the Research Ethics Committee. Our research is compliant with the HFEA Code of Practice and has undergone multiple inspections by the HFEA since the licence was granted.

Informed consent was obtained from all couples that donated spare embryos following infertility treatment. Before giving consent, donors were provided with information about the research project, an opportunity to receive counselling and the conditions that apply to the research licence. No financial inducements were offered for donation. Embryos surplus to the patient’s IVF treatment were donated cryopreserved and were transferred to the University of Cambridge and Francis Crick Institute, where they were thawed and used in the research project.

### Mouse embryo collection

Female mice, aged four to eight weeks (C57BL6 × CBA) F1, were superovulated by intraperitoneal injection of 5 IU of pregnant mare serum gonadotrophin (PMSG; Sigma-Aldrich) followed 48hours later with an intraperitoneal injection of 5 IU of human chorionic gonadotrophin (HCG; Sigma-Aldrich) and mating with eight weeks or older (C57BL6 × CBA) F1 males. The mice were maintained under a 12-hour light–dark cycle. Zygotes were isolated from oviducts of plugged mice on at 0.5 days post fertilisation in FHM medium (Merck; MR-122-D) under mineral oil (Origio; ART-4008-5P), and cumulus cells were removed using hyaluronidase (Sigma-Aldrich; H4272). All procedures involving animals were conducted in accordance with the UK Home Office Licence Number PP8826065.

### Viral transduction

Mouse embryos were transduced with Cellight™ Histone 2B-GFP (Cat. No. C10594). Live embryo imaging was conducted after 24hr using a confocal microscope. For AAV6 transduction, embryos were co-cultured with scAAV6-tdTomato (Plasmid #59462) at varying concentrations: 1 × 10×8 IU/mL, 1 × 10×10 IU/mL, 1 × 10×12 IU/mL, or without AAV6, for 24 hours. Subsequently, the embryos were cultured *in vitro*, and the expression of tdTomato was assessed after 24 hrs and 48hrs using fluorescence microscopy. For lentivirus transduction, mouse embryos were transduced with a high-titer lentivirus carrying an H2B-GFP reporter (Addgene plasmid #26777) and imaged after 24hrs using a confocal microscope.

### Live embryo staining

Mouse embryos were cultured in KSOM media (Merk; MR-101-D) supplemented with various dyes at working concentrations: SPY650-DNA (1:1,000, Spirochrome), Nuclight Rapid Red (1:1,000, Incucyte #4717), 1 μM of 5-TMR-Hoechst, 1 μM of 4-TMR-Hoechst, 1 μM of 4-580CP-Hoechst, and 1 μM of 5-580CP-Hoechst^52^ (gift from Gražvydas Lukinavičius lab) and were subsequently imaged after 48hrs.

### Generation of modified mRNAs by *in vitro* transcription

mRNA synthesis was performed as previously described ^48^. The dsDNA template (H2B-mCherry Plasmid 20972 from Addgene) was linearized, and a small aliquot of the digestion mix was subjected to gel electrophoresis to verify complete digestion. Linearised plasmid was purified using a PCR purification kit (Qiagen, Cat. No. 28104). Poly(A) tailing was carried out using KAPA PCR ready mix (2X) and the following primer sets: the forward primer

(CTTACTGGCTTATCGAAATTAATACGA) and the reverse primer (TTTTTTTTTTTTTTTTTTTTTTTTTTTTTTTTTTTTTTTTTTTTTTTTTTTTTTTTTTTTT TTTTTTTTTTTTTTTTTTTTTTTTTTTTTTTTTTTTTTTTTTTTTTTTTTTTTTTTTTTTTTTTTTTAAACAACAGATGGCTGGCAACTAGAAGG) from Integrated DNA Technologies. Subsequently, the digested plasmid was adjusted to a concentration of 10 ng/μl. Tail PCR was run for 32 cycles and purified using PCR purification kit. In vitro transcription was performed using MEGAscript T7 kit (Thermo Fisher; Cat. No. AMB13345): custom NTP mix was prepared with 3’-O-Me-m7G cap analogue (60 mM, NEB), GTP (75mM, MEGAscript T7 kit), ATP (75mM, MEGAscript T7 kit), Me-CTP (100 mM, TriLink; Cat. No. N-1014-1) and pseudo-UTP (100mM, Tri-link; Cat. No. O-0263). Reaction was heated at 37 C for 2 h. 2 ul of Turbo DNase (Thermo Fisher; Cat. No. AM2238) was added and incubated at 37 C for 15 min. DNAse treated reaction mix was purified using RNAeasy kit (Qiagen; Cat. No. 74104) according to the manufacturer’s instructions. RNA was phosphatase-treated using Antarctic phosphatase (New England BioLabs; Cat. No. M0289S) and purified using MEGAclear kit (Thermo Fisher, Cat. No. AM1909).

### Human and mouse embryo culture

Slow frozen human embryos were thawed using the Blast thaw kit (Origio; Cat No. 10542010A), following the manufacture’s instructions. Vitrified human embryos were thawed using Vit Kit-Thaw (Fujifilm; Cat No. 90137-SO). Mouse embryos and human embryos were cultured at 37 °C and 6% CO2 in drops of pre-equilibrated Global medium (LifeGlobal; Cat No. LGGG-20) supplemented with 5 mg/ml protein supplement (LifeGlobal; Cat No. LGPS-605) and covered with mineral oil (Origio; Cat No. ART-4008-5P).

### Immunofluorescence and confocal imaging

Embryos were fixed using freshly prepared 4% paraformaldehyde in PBS either at room temperature for 1 hr or overnight at 4 °C, followed by three washes in 1X PBS with 0.1% Tween-20 (Sigma-Aldrich; P1379-25ML) to eliminate residual paraformaldehyde. Subsequently, embryos were permeabilized with 1X PBS with 0.5% Triton X-100 and then placed in blocking solution (3% BSA (Sigma-Aldrich; 05470-5G) in 1X PBS with 0.2% Triton X-100 (Sigma-Aldrich X100-5ML)) for 2 h at room temperature on a rotating shaker. Embryos were then incubated overnight at 4°C on rotating shaker with primary antibodies diluted in blocking solution, at the following concentrations: CDX2 (BioGenex; MU392A-UC) at 1:20, NANOG (2B Scientific, REC-RCAB0001P) at 1:100 and GATA3 (Abcam, ab199428) at dilution 1:100. The following day, embryos were washed in 1X PBS with 0.2% Triton X-100 for 20 min at RT on a rotating shaker and then incubated with secondary antibodies diluted in blocking solution for 1 h at room temperature on a rotating shaker in the dark. Next, embryos were washed in 1X PBS with 0.2% Triton X-100 for 20 min at room temperature on a rotating shaker. Finally, embryos were placed in 1X PBS with 0.1% Tween-20 with Vectashield and DAPI mounting medium (Vector Lab; H-1200) (1:30 dilution). Embryos were placed on µ-Slide 8 well dishes (Ibidi; 80826) for confocal imaging. Confocal immunofluorescence images were taken with SP8 confocal microscope (Leica Microsystems) and 2-μm-thick optical sections were collected.

### Electroporation

Mouse and human embryos were washed and placed in drops of Opti-MEM (#31985062, Thermo Fisher Scientific). The dish was then placed on a heated microscope stage (Olympus IX70). We transferred 7 µL of the H2B-mCherry mRNA solution onto an electroporation chamber between the electrodes of the plate (CUY501P1-1.5, NEPA GENE Co. Ltd, Sonidel, Ireland). The impedance was adjusted to between 0.19Ω and 0.21Ω (typically 0.20 Ω) by either adding or removing the electroporation solution. The electroporation parameters of mouse embryos were as follows: 1 poring pulses of 0.1 V, lasting 50 ms with a 50 ms interval, 10% decay, immediately followed by 2 transfer pulses of 20 V, 40% decay, lasting 25 ms with a 50-ms interval. The electroporation parameters for human embryos were as follows: 6 poring pulses of 15 V, lasting 2 ms with a 50 ms interval, 10% decay, immediately followed by 5 transfer pulses of 5 V, 40% decay, lasting 50 ms with a 50 ms interval. Immediately after electroporation, the embryos were removed from the electroporation chamber, washed and cultured in equilibrated Global medium (#LGGG-20; LifeGlobal) supplemented with 5 mg/ml protein supplement (#GHSA125: LifeGlobal) and covered with mineral oil.

### Live embryo imaging

Electroporated embryos were placed in a fluorinated ethylene propylene (FEP) foil microwell sample holder, containing equilibrated Global medium (LifeGlobal; LGGG-20), supplemented with 5 mg/ml protein supplement (LifeGlobal; LGPS-605), and covered with mineral oil (Origio; ART-4008-5P). The Viventis LS1 and LS2-Live dual illumination and inverted detection microscopes were used for live imaging embryos. Time-lapse images of embryos were captured every 15 minutes for up to 2 days at 37°C and 6% CO2 with either a 10× 0.3 NA or a 25× 1.1 NA. Volumes of 400-500 μm were acquired with a Z spacing of 2 μm between slices and a 100 ms exposure time for each slice. Laser intensity was minimized to obtain a reasonable signal-to-noise ratio from the raw data while minimizing phototoxicity. The reconstruction of movies was performed using IMARIS software (Bitplane, AG) and Fiji ImageJ open-source image processing package.

### 3D nuclear segmentation and tracking

Semi-automated 3D nuclear segmentation and tracking was performed using 1) a custom version (https://github.com/akarsa/anisotropic_stardist_3d) of the StarDist-3D network ^49^ tailored to our data, and 2) cutting-edge, regularized Gaussian mixture optimal transport ^53^.

Both softwares are available for download (https://github.com/akarsa/star-3d, https://github.com/akarsa/optimal3dtracks).

As a first pre-processing step, averaging of dual-view light-sheet images was carried out, where available, for contrast enhancement. Furthermore, to avoid over-segmentation and account for variations in imaging parameters, we resampled the data to an optimized 0.23 µm lateral and 2.67 µm axial resolution, normalized the intensity, and applied gamma correction. Samples suffering from more significant axial degradation were further down sampled axially. Model-based deconvolution ^54^ was also used as an alternative. The StarDist3D network architecture was altered to increase the receptive field of the network, thus being able to cope with the increased size and shape variability in (human) embryo data and trained this on annotated mouse embryo images (https://github.com/akarsa/star-3d).

Nuclei were automatically segmented over 3D timelapses of 100 time points. Besides nuclei, the custom StarDist-3D network also delineates various unwanted, nuclei-like structures such as dirt in and around the embryo. As a pre-filtering step, features smaller than 30 µm3 in size were removed.

The segmented nuclei were tracked using regularized Gaussian mixture optimal transport (GMMOT) (https://github.com/akarsa/optimal3dtracks, https://github.com/judelo/gmmot, ^53^ to calculate transition probabilities between positions at consecutive time points. A regularization term was added to GMMOT to improve the quality of the automated tracking.

Using an in-house Python script, the automatically generated 3D segmentations and tracks were converted to a format readable by Fiji’s TrackMate tool ^55^. An expert biologist (A.A.) curated and corrected the results in TrackMate.

Videos of 2D axial projections of both the intensity images and segmentations are available in the supplementary materials; tracks were indicated by preservation of label colour over time. The intensity images were masked to better visualise the region of interest (embryo). The videos also indicate the evolutions of the dendograms, and number of nuclei detected in the embryo. Several metrics were calculated to evaluate and correlate the sizes, shapes, and orientations of the embryo and the segmented nuclei. For each segmented nuclei, the volume, anisotropy (computed as ratio between longest axis diameter and shortest axis diameter of the best fitting ellipsoid describing the nuclei), centroid, and orientation (3D direction of the long axis) were calculated. The same statistics were computed for the entire embryo, i.e. the convex hull of all segmented nuclei, at each time point. From these metrics, the “average shell thickness” (standard deviation of distance between the nuclei and the embryo center), angle between the embryo and nuclei orientations (α), nuclei orientation with respect to the embryo center (β), and cell division direction (ϑ) with respect to the embryo center^19^ could also be computed. Linear regression analysis was used to investigate correlations between different measures, and Kolmogorov–Smirnov test was performed to statistically evaluate the deviations of β and ϑ from the uniform distribution.

### Quantifications and statistical analysis

Statistical details for each experiment are described in the corresponding figure legends. Biological replicates are denoted as ‘n’. For microscopy data, Graphpad Prism software was used to perform statistical analyses, using the Mann-Whitney U test or unpaired, two-tailed t-test. Values are presented as mean ± SD. Statistical differences are represented as follows: * p < 0.05, ** p < 0.01, *** p < 0.001.

## Acknowledgments

We thank the donors of human embryos whose contributions were essential for this research. We thank all members of the Niakan lab for their technical assistance, advice and comments on the manuscripts. We thank the Centre for Trophoblast Research for technical support and advice. We thank Florian Holfelder and Timo Kohler for support with confocal microscopy. We thank the Francis Crick Institute’s Advanced Light Microscopy Science Technology members (Kurt Anderson, Alessandro Ciccarelli, Camille Charoy, and Todd Fallesen) for imaging advice and assistance. We thank Molly Strom at the Francis Crick Institute’s Viral Vector Service for producing viral vectors used in this project. We thank Gražvydas Lukinavičius lab for providing us with live DNA dyes (5-TMR-Hoechst, 4-TMR-Hoechst, 4-580CP-Hoechst, and 5-580CP-Hoechst). We also would like to thank Mary Herbert, Peter Rugg-Gunn and Azim Surani for their valuable feedback on our study. Work in the laboratory of KKN was supported by the Wellcome 221856/Z/20/Z and the Wellcome Human Developmental Biology Initiative 215116/Z/18/Z. Work in the laboratory of KKN was also supported by the Francis Crick Institute which receives its core funding from Cancer Research UK FC001120, the Medical Research Council FC001120 and Wellcome FC001120. AK and LM were supported by EPSRC EP/Y008715/1. For the purpose of Open Access, the authors have applied a CC BY public copyright licence to any Author Accepted Manuscript version arising from this submission.

## Author contribution

AA and KKN conceived the project, interpreted the data, and wrote the original draft of the manuscript. AA designed and performed experiments and generated figures. AM developed the mouse and human embryo electroporation methods. AK and LM developed the image segmentation and lineage tracking analysis pipeline, generated figures and interpreted data. AM did all the human embryo thaws and electroporation at the Francis Crick Institute. AA and KKN did all the human embryo thaws and electroporation at the University of Cambridge. IL and BJS assisted with setup of light sheet microscopy and interpretation of the data. KE, PS, LC, and MW coordinated human embryo consenting and facilitated donations.

## Declaration of interests

The authors declare no competing interests.

## Data availability statement

Source data are provided with this paper. All other data supporting the findings of this study are available from the corresponding author upon reasonable request. The primary microscopy data will be uploaded at publication.

## Code availability

Semi-automated 3D nuclear segmentation and tracking was performed using 1) a custom version (https://github.com/akarsa/anisotropic_stardist_3d) of the StarDist-3D network tailored to our data, and 2) cutting-edge, regularized Gaussian mixture optimal transport. Both are available for download (https://github.com/akarsa/anisotropic_stardist_3d, https://github.com/akarsa/cell_tracking_with_optimal_transport).

The StarDist3D network architecture was altered to increase the receptive field of the network, thus being able to cope with the increased size and shape variability in (human) embryo data and trained this on annotated mouse embryo images (https://github.com/akarsa/anisotropic_stardist_3d).

The segmented nuclei were tracked using regularized Gaussian mixture optimal transport (GMMOT) (https://github.com/akarsa/cell_tracking_with_optimal_transport, https://github.com/judelo/gmmot, to calculate transition probabilities between positions at consecutive time points.

## Supplementary Data

**Extended Data Fig 1.**
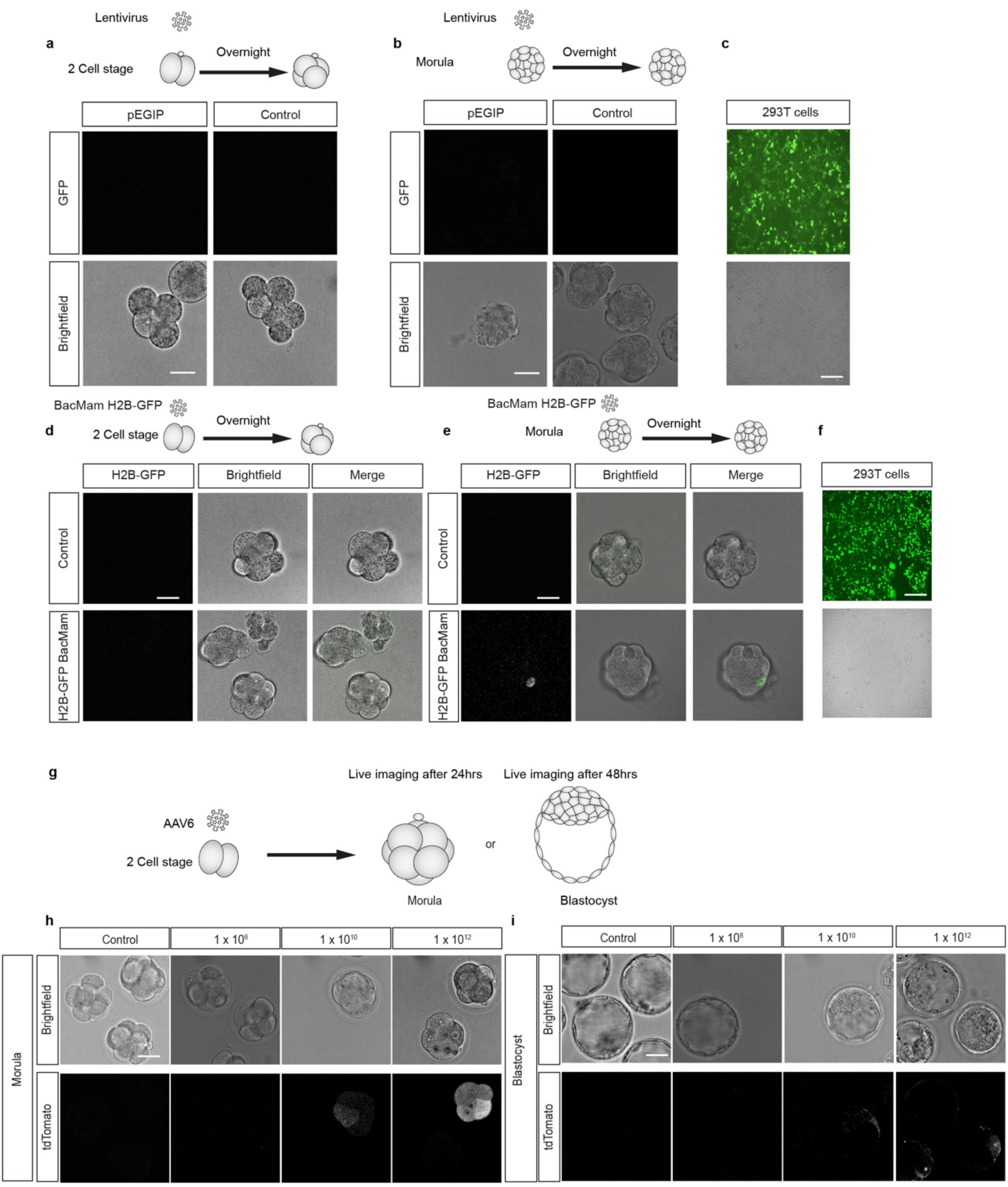
Transduction of early mouse embryos and HEK 293 T cells with lentiviral, baculovirus mammalian expression system, and adeno-associated virus vectors. **a,b** Representative images of mouse embryos transduced with a pEGIP lentiviral vector that carries an EGFP gene downstream of the EF-1a promoter. Embryos were washed tree times with hyaluronidase for 10 seconds and cultured in global media. The zona pellucida of mouse embryos was removed prior to transduction. Mouse zona-free embryos were then divided into two groups for transduction at the: **(a)** two cell stage or at the **(b)** morula stage. Embryos were culture in Global medium containing pEGIP lentiviral vectors or in control Global medium in the absence of lentivirus. Embryos were then imaged live 24 hours following transduction. **c,** Transduction of HEK 293 T cells with concentrated pEGIP lentivirus was performed as a control confirming efficacy of EGFP lentiviral vectors. **d,e** CellLight™ Histone H2B-GFP, BacMam 2.0 baculovirus transduction of early mouse embryos from the **(d)** 2-cell stage or the **(e)** morula stage. H2B-GFP BacMam transduced mouse embryos were imaged 24 hrs after transduction. **f,** Transduction of HEK 293 T cells with H2B-GFP BacMam was performed as a control confirming GFP expression. **g,** Schematic of AAV6-mediated transduction of embryos. **h,i** Mouse 2-cell stage embryos were cultured in the presence of AAV6 adeno-associated virus carrying a tdTomato gene downstream of the EF-1a promoter. The expression of tdTomato was analyzed by live cell imaging at the **(h)** morula or **(i)** blastocyst stage. Scale bar 30µm.

**Extended Data Fig 2.**
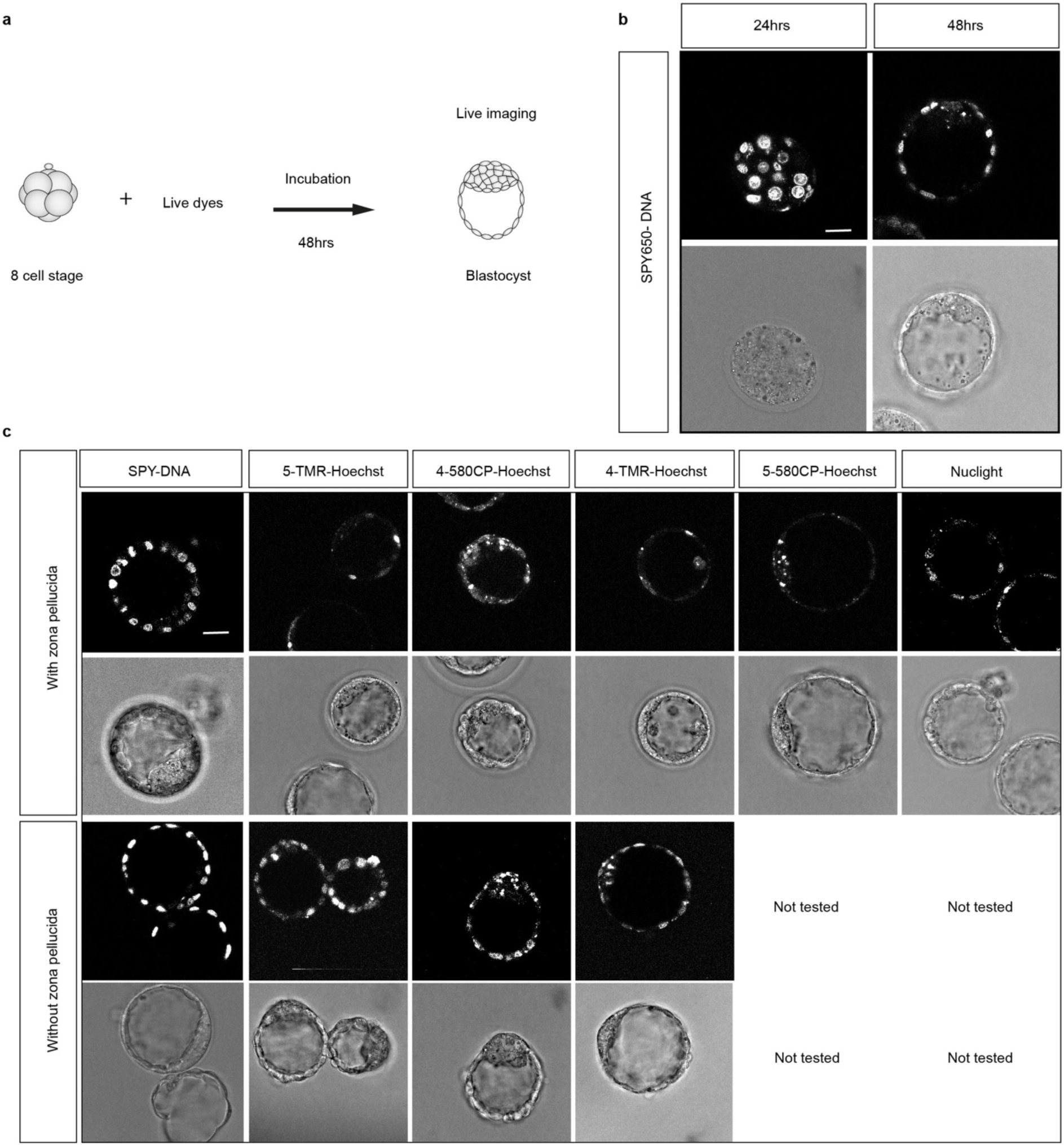
Staining of mouse embryos followed by live-cell imaging using different live DNA staining dyes. **a,** Schematic representation of 8-cell stage mouse embryos cultured in Global medium containing live dyes for 24 or 48 hrs until the blastocyst stage. Mouse embryos were subsequently imaged on a confocal microscope in the presence of the dyes. **b,** Representative images of embryos stained with SPY650-DNA after 24 hrs and 48 hrs and phase-contrast images. Embryos were stained from the 8-cell stage until the times indicated. **c,** Representative images of embryos stained with various live DNA dyes (SPY650-DNA, 5-TMR-Hoechst, 4-TMR-Hoechst, 4-580CP-Hoechst, 5-580CP-Hoechst, and NucLight Rapid Red). Imaging was conducted 48 hrs post-staining. Phase contrast images are shown. Scale bar, 30 μm.

**Extended Data Fig 3.**
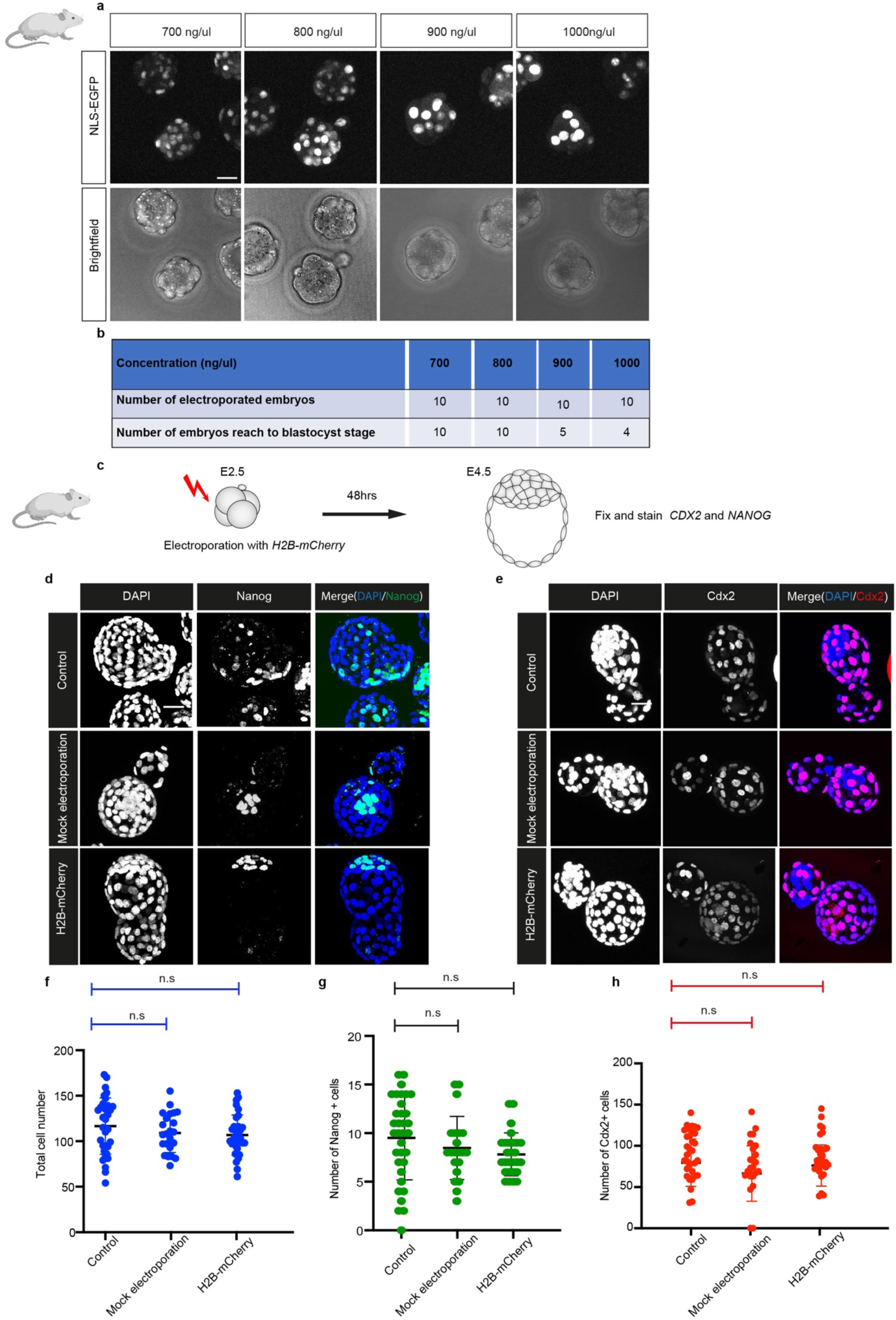
Development of mouse embryos electroporated with various concentrations of mRNA. **a,** Mouse embryos were electroporated at 4-cell stage with an in vitro transcribed EGFP mRNA containing a nuclear localisation signal (NLS) at the concentrations shown. Fluorescent and phase-contrast images were taken 24 hrs after electroporation. **b,** The number of embryos developing to the blastocyst stage is summarised. n = 10 embryos electroporated per concentration. **c,** Schematic of the experimental setup of mouse embryos electroporated with 500ng/µl H2B-mCherry mRNA followed by 48 hrs culture in Global media to the blastocyst stage. **d,e** Control mouse embryos cultured without electroporation (n= 33), a mock electroporation group (n=23), and embryos electroporated with 500ng/µl H2B mCherry mRNA (n=32) were immunofluorescently analysed for the expression of **(d)** Nanog (epiblast molecular marker), **(e)** CDX2 (trophectoderm molecular marker) and DAPI nuclear staining. **f,** Quantification of total cell number; **g,** NANOG-positive cells; or **h,** CDX2-positive cells. No difference in the number of Cdx2-positive and Nanog-positive cells was observed. P > 0.05 NS, not significant, Student’s t-test. Scale bars, 30 μm. Error bars represent s.d.

**Extended Data Fig 4.**
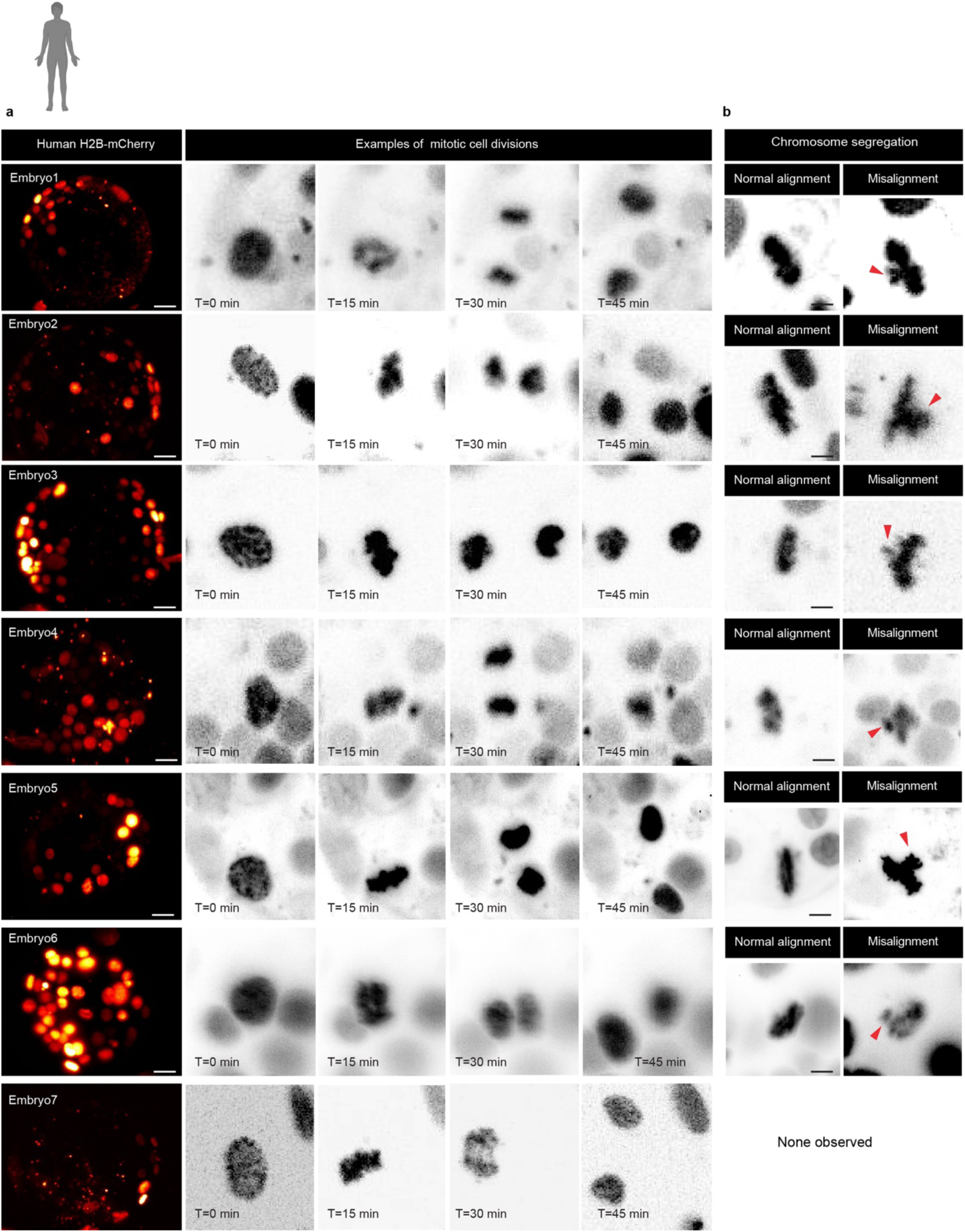
Live imaging of human embryos labelled with H2B-mCherry using light-sheet microscopy to visualize mitosis. **a,** Time-lapse images of mitosis (prophase, metaphase, anaphase and telophase) in human cells at the blastocyst stage. Human embryos were imaged following H2B-mCherry nuclear labelling (n = 7). **b,** Examples of normal alignment and misalignment of chromosomes observed in human blastocyst cells. Misalignment indicated with an arrowhead.. Scale bar, 30μm.

**Extended Data Fig 5.**
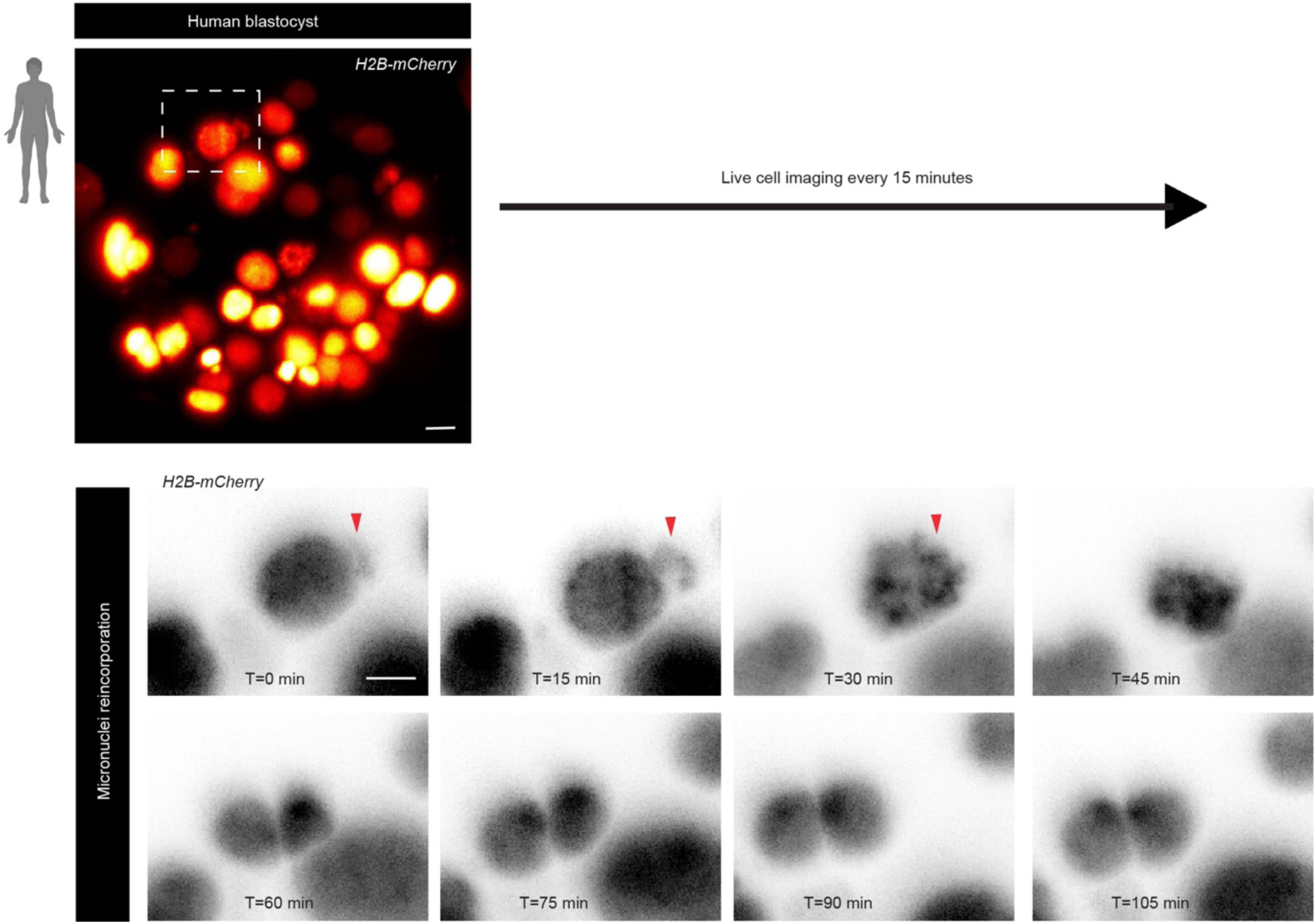
Micronuclei reincorporation into the main nucleus during mitosis. Example of micronuclei reincorporating during mitosis in a human embryo expressing H2B-mCherry. Micronuclei indicated with an arrowhead. Scale bars, 15 μm.

**Extended Data Fig 6.**
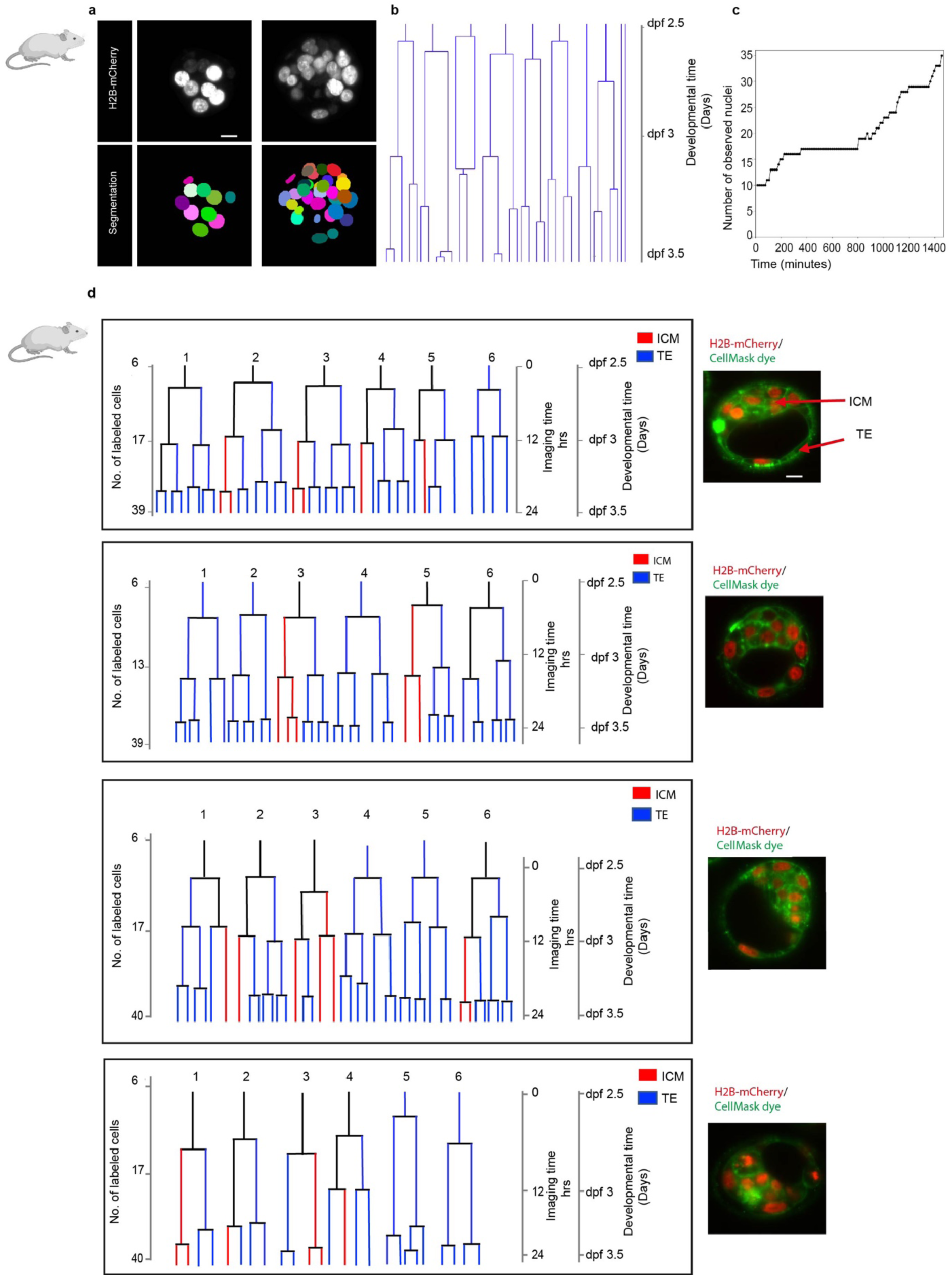
Fate of cells in mouse preimplantation embryos. **a,b,** An example of 3D segmentation and tracking of mouse embryo nuclei following H2B-mCherry mRNA electroporation. **c,** Number of observed nuclei over time. **d,** Lineage trees of mouse embryos expressing H2B-mCherry from the 8-cell to blastocyst stage following light-sheet live embryo imaging. ICM (red) or TE (blue) cells were assigned based on position and is displayed on the tree by color code. Representative z-section fluorescent image of the mouse embryos is shown for both H2B-mCherry and CellMask membrane dye. Scale bars, 15 μm.

**Extended Data Fig 7.**
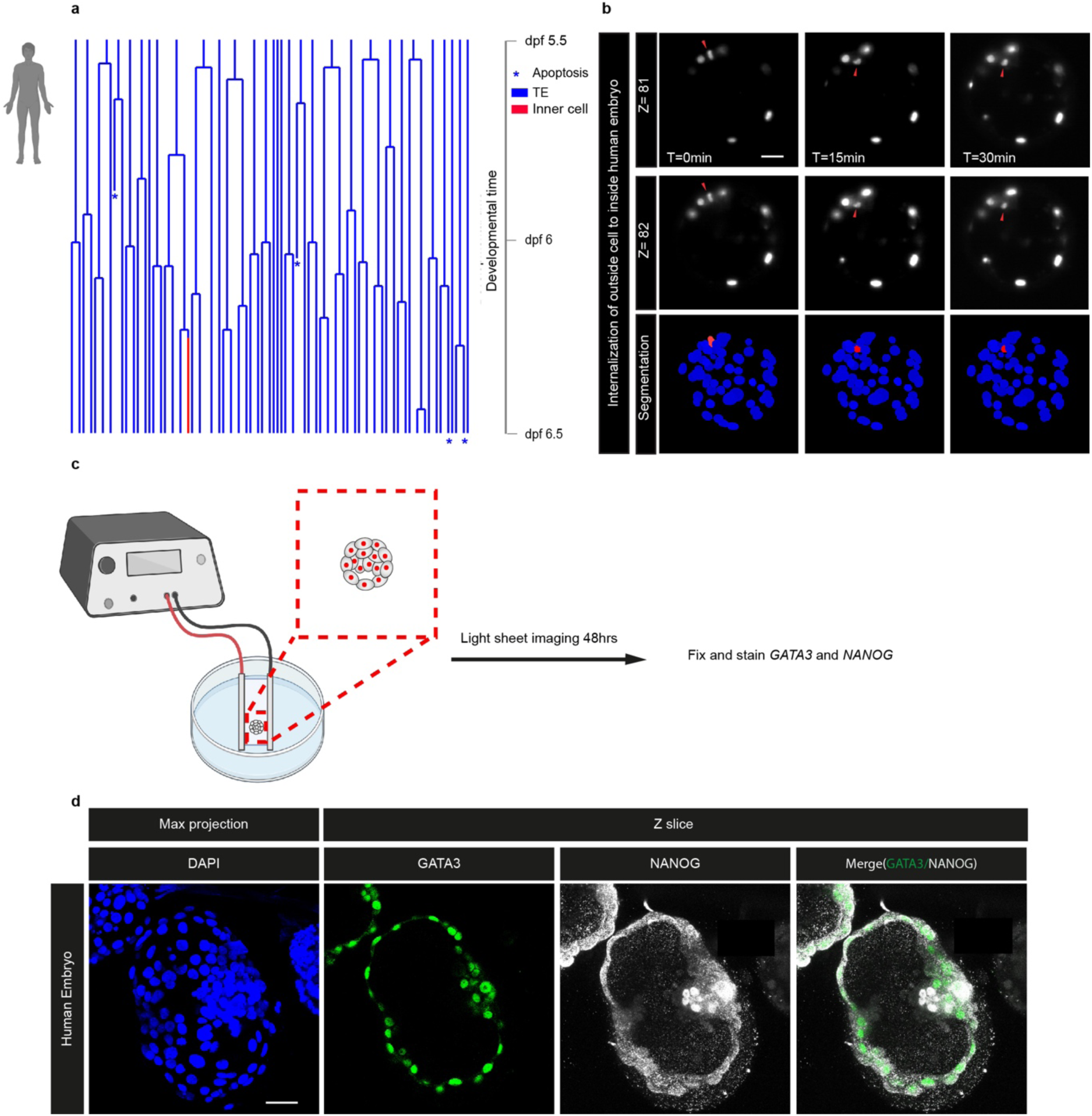
Fate of TE cells in human embryos from early to late blastocyst stages. **a,** Lineage tree of H2B-mCherry labelled nuclei of human pre-implantation embryo from the early to late blastocyst stage. Each cell represented on the tree is color coded according to position. **b,** Selected frames from time-lapse imaging of an H2B-mCherry expressing human embryo. During the blastocyst stage, a transient-outer cell expressing H2B-mCherry was observed to migrate inward. The nucleus, marked with red arrow, signifies a transient outer cell moving inward during the late blastocyst stage. **c, d,** human embryo electroporated with 500ng/µl H2B mCherry mRNA were immunofluorescently analysed for the expression of NANOG (epiblast molecular marker), GATA3 (trophectoderm molecular marker) and DAPI nuclear staining. Scale bar: 30 μm. See Video S7, S8, S9, S10, S11.

**Extended Data Fig 8.**
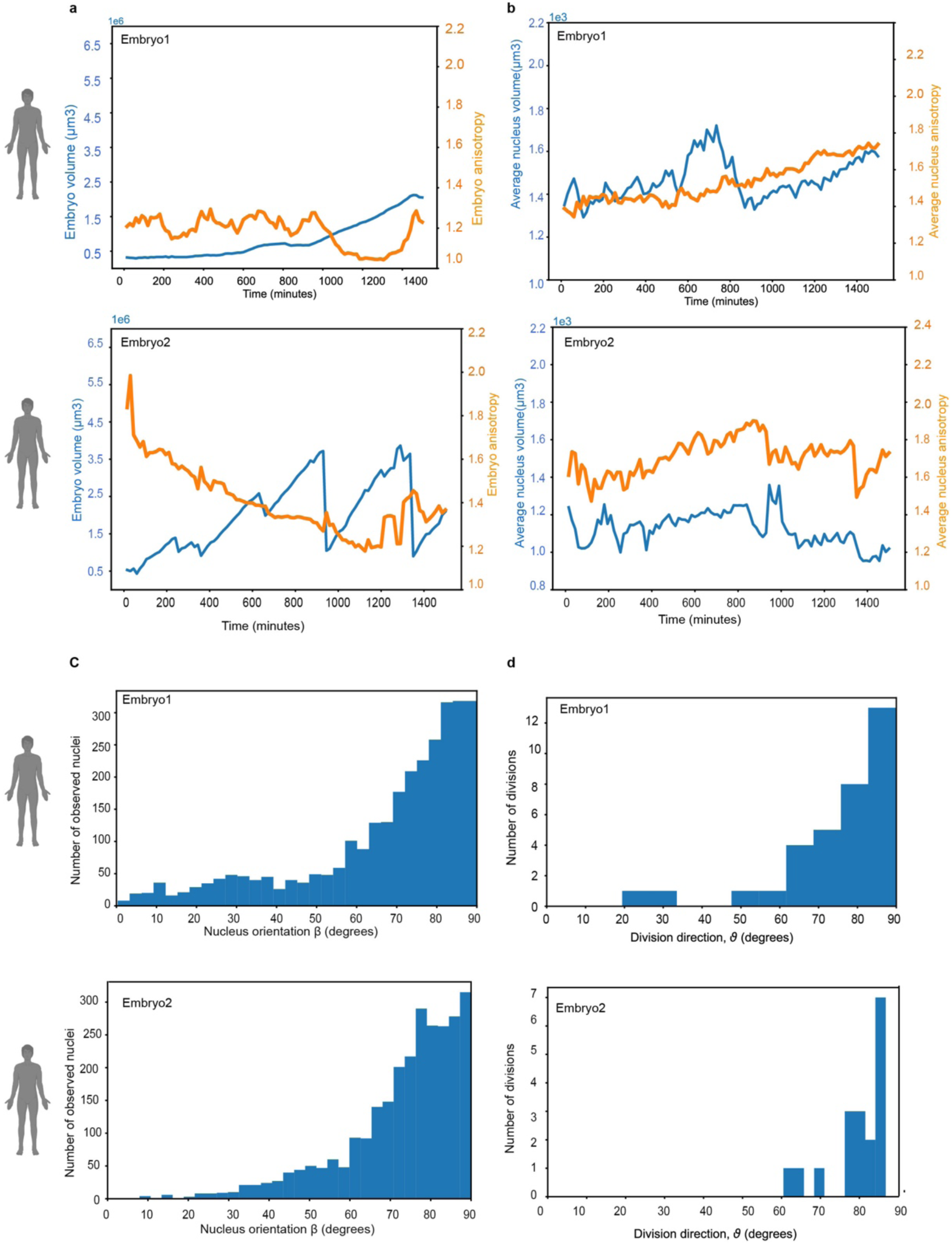
Nuclei shape, size, and division axis orientation of cells in human blastocysts. **a,** Human embryos volume (blue) and anisotropy (orange) measured across time. **b,** Average nucleus volume (blue) and anisotropy (orange) measured across time. **c,** Histogram of the angle β of the nucleus orientation across all time points showing that nuclei prefer to be oriented tangentially to the embryo center rather than radially oriented (K-S test p<10^−300^). **d,** histogram of ϑ across all time points showing that cells strongly prefer to divide tangentially rather than radially to the centre of the embryo (K-S test p∼10^−12^).

## Supplementary Videos 1-11

**Supplementary video 1: Live time-lapse imaging of mouse pre-implantation development.** Mouse embryos were electroporated at the 8-cell stage with H2B-mCherry mRNA. Fluorescent and phase-contrast images were taken for 48 hours. The movie displays maximum projections of live-cell time-lapse imaging of mouse pre-implantation embryos expressing nuclear H2B-mCherry.

**Supplementary video 2: Live time-lapse imaging of human blastocyst development labeled with H2B-mCherry.** Human blastocysts were electroporated with H2B-mCherry mRNA. Fluorescent images were taken every 15 minutes for 46 hours.

**Supplementary video 3: Examples of lagging chromosomes observed in mouse and human blastocyst cells.**

**Supplementary video 4: Examples of mitotic slippage observed in mouse and human blastocyst cells.**

**Supplementary video 5: Examples of normal and multipolar cell division observed in human blastocyst cells.**

**Supplementary video 6: Examples of micronuclei inheritance observed in mouse and human blastocyst cells.**

**Supplementary video 7: Fate of TE cells in human embryo 1 from early to late blastocyst stages.** The movie displays segmentation of human embryo nuclei labeled with H2B-mCherry and complete lineage tree from the early to late blastocyst stage, as well as the number of observed nuclei across time.

**Supplementary video 8: Fate of TE cells in human embryo 2 from early to late blastocyst stages.** The movie displays segmentation of human embryo nuclei labeled with H2B-mCherry and complete lineage tree from the early to late blastocyst stage, as well as the number of observed nuclei across time.

**Supplementary video 9: Fate of TE cells in human embryo 3 from early to late blastocyst stages.** The movie displays segmentation of human embryo nuclei labeled with H2B-mCherry and complete lineage tree from the early to late blastocyst stage, as well as the number of observed nuclei across time.

**Supplementary video 10: Demonstration of transient-outer cell to inside human blastocyst.** The movie displays a 3D view of a mother cell dividing into two daughter cells, with one remaining outside and the other undergoing the internalization process. The final position of the daughter cells is shown. The nucleus of the mother cell and the daughter cells are marked with green spots. Arrows mark the final positions of the daughter cells.

**Supplementary video 11:** The movie displays segmentation of transient-outer cell moving inwards. The transient outer daughter cell is marked with red color.

## Notes

### Competing Interest Statement

The authors have declared no competing interest.

